# A hierarchical clustering and data fusion approach for disease subtype discovery

**DOI:** 10.1101/2020.01.16.909382

**Authors:** Bastian Pfeifer, Michael G. Schimek

## Abstract

Recent advances in multi-omics clustering methods enable a more fine-tuned separation of cancer patients into clinical relevant clusters. These advancements have the potential to provide a deeper understanding of cancer progression and may facilitate the treatment of cancer patients. Here, we present a simple hierarchical clustering and data fusion approach, named *HC-fused*, for the detection of disease subtypes. Unlike other methods, the proposed approach naturally reports on the individual contribution of each single-omic to the data fusion process. We perform multi-view simulations with *disjoint* and *disjunct* cluster elements across the views to highlight fundamentally different data integration behaviour of various *state-of-the-art* methods. *HC-fused* combines the strengths of some recently published methods and shows superior performance on real world cancer data from the TCGA (The Cancer Genome Atlas) database. An R implementation of our method is available on GitHub (*pievos101/HC-fused*).

## 1 Introduction

The analysis of multi-omic data has great potential to improve disease subtyping of cancer patients and may facilitate personalized treatment [1, 2]. While single-omic studies have been conducted extensively in the last years, multi-omics approaches, taking into account data from different biological layers, may reveal more fine-grained insights on the systems-level [3–5]. However, the analysis of data sets from different sources like DNA sequences, RNA expression, and DNA methylation brings great challenges to the computational biology community. One of the major goals in integrative analysis is to cluster patients based on features from different biological layers to identify disease subtypes with enriched clinical parameters. Integrative clustering can be divided into two groups. Horizontal integration on the one side, which is the aggregation of the same type of data, and vertical integration on the other side, which concerns the analysis of heterogeneous omics data sets for the same group of patients [6]. In addition to this classification, one distinguishes between early and late integration approaches. Late integration-based methods first analyze each omics data set separately and then concatenate the information of interest to a global view. Early integration first concatenates the data sets and then performs the data analysis.

In vertical integration tasks, one major problem is that the data sets are often highly diverse with regard to their probabilistic distributions. Thus, simply concatenating them and applying single-omics tools is most likely to bias the results. Another issue arises when the number of features differs across the data sets with the effect that more importance is assigned to a specific single-omics input. The recent years have seen a wide range of methods that aim to tackle some of these problems [7]. Most prominent is *SNF* (*Similar Network Fusion*) [8]. For each data type it models the similarity between patients as a network and then fuses these networks via an interchanging diffusion process. Spectral clustering is applied to the fused network to infer the final cluster assignments. A method which builds up on *SNF* is called *NEMO* and was recently introduced in [9]. This paper provides solutions to partial data and implements a novel *eigen-gap* method [10] to infer the optimal number of clusters. A method called *rMKL-LPP* [11] makes use of dimension reduction via multiple kernel learning [12] in order to perform the data integration step. In addition, overfitting is taken care of by regularization.

*Spectrum* [13] is another recently published multi-omics clustering method and R-package. Again it performs spectral clustering but provides a data integration method which is significantly different from *NEMO* and *SNF. Spectrum*’s data integration approach is based on a tensor product graph (TPG) diffusion technique. Furthermore, it provides a novel method to infer the optimal number of clusters *k* using eigenvector distribution analysis. Statistical solutions for the clustering of heterogeneous data sets were introduced in [14]. In the (*iClusterPlus*) approach the observations from different genomic data types are simultaneously regressed under appropriate distributional assumptions to a common set of latent variables [14]. A computational intensive Monte-Carlo Newton-Raphson algorithm is used to estimate the parameters. Also, a fully Bayesian version of *iClusterPlus* was recently put forward in [15]. A number of additional techniques have been developed as outlined in [16]. One of these techniques is called *PINSPlus* [17, 18]. Its authors suggest to systematically add noise to the data and to infer the best number of clusters based on the stability against this noise. When the best *k* (number of clusters) is detected, binary matrices are formulated reflecting the cluster solutions for each *single-omics*. A final agreement matrix is derived by counting the number of times two patients appear in the same cluster. This agreement matrix is than used for a standard clustering method, such as *kmeans*.

In this article we introduce a new method for hierarchical data fusion and integrative clustering called *HC-fused*. First we cluster each data type with a standard hierarchical clustering algorithm. We then form network structured views of the omics data sets, and finally apply a novel approach to combine these views via a hierarchical data fusion technique. In contrast to other methods, *HC-fused* naturally reports on the contribution of the individual views to the data fusion process. Its advantage is the adoption of simple data analytic concepts with the consequence that results can be easily followed-up and interpreted.

## 2 Materials and methods

### 2.1 Data preprocessing

Data normalization and imputation is done as suggested by [8]. When a patient has more than 20% missing values we do not consider this patient for further investigation. When a specific feature has more than 20% missing values across the patients, we remove this feature from further investigation. The remaining missing values are imputed with the *k*-Nearest Neighbor method (*k*NN). Finally, the normalization is performed as folllows:

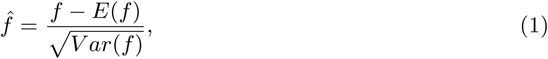

where *f* is a biological feature.

### 2.2 Transforming the views into network structured data

Given is a set of data views 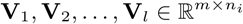, where *m* is the number of observations, and *n*_*i*_ is the number of features from view **V**_*i*_. We transform these views into connectivity matrices **G**_1_, **G**_2_, …, **G**_*l*_ ∈ {0, 1} ^*m*×*m*^. This is done by clustering the views with a hierarchical clustering algorithm using Ward’s method [19]. We then infer the best number of clusters *k* via the *Silhouette Coefficient*. The produced matrices **G**_1_, **G**_2_, …, **G**_*l*_ are binary matrices with entry 1 when two elements are connected (being in the same cluster), and 0 otherwise. In addition, we construct a **G**_∧_ matrix as follows:

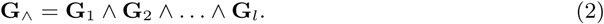

The matrix **G**_∧_ reflects the connectivity between patients confirmed by all views.

### 2.3 Generating the fused similarity matrix P

For data fusion we apply a bottom-up hierarchical clustering approach to the binary matrices **G**. Initially, no patient cluster exists. In each iteration, patients or clusters of patients (*c*_1_ ∈ *C* and *c*_2_ ∈ *C*) fuse to a newly built cluster with minimal distance *d*_*min*_, until just one single cluster remains. The distance between two clusters is calculated as

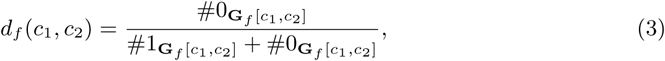

where *f* ∈ {1, …, *l*, ∧} and # means count.

In each iteration the algorithm is allowed to use the distances from any given binary matrix **G**_1_, **G**_2_, …, **G**_*l*_, **G**_∧_. We refer to **G**_*min*_ as the matrix containing the minimal distance *d*_*min*_, where *min* ∈ 1 … *l*, ∧. In cases where the minimal distance is shared by multiple matrices we give preference to fusing the clusters in **G**_∧_. In our approach, a fusion event between two clusters is denoted as (*c*_1_ ++ *c*_2_) or *fuse*(*c*_1_, *c*_2_). During the fusion process we count how many times a patient pair (*i, j*) occurs in the same cluster. This information is captured in the fused similarity matrix **P** ∈ ℝ^*m*×*m*^.

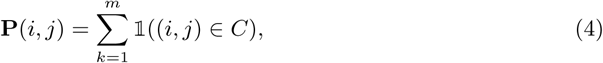

where 𝟙 denotes the indicator function.

Finally, the matrix **P** is normalized by its maximal entry. The similarity matrix **P** can be used as input for arbitrary cluster algorithms. Currently we apply agglomerative hierarchical clustering using Ward’s method [19] as implemented in [20].

### 2.4 Contribution of the views to the data fusion operation

We define matrices **S**_1_, **S**_2_, …, **S**_*l*_, **S**_∧_ ∈ ℝ^*m*×*m*^ providing information about the contribution of each view to the data fusion process. We count how many times a patient is involved in a fusion operation (*c*_1_ ++ *c*_2_) and in what view this fusion is executed. This information is captured in the source matrices **S**.

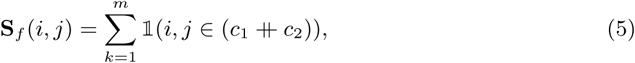

where *f* ∈ {1, …, *l*, ∧} and 𝟙 is the indicator function. The clusters *c*_1_ and *c*_2_ refer to the clusters which are fused in iteration *k*.

It should be noted, however, that in each fusion operation there might occur multiple minimal distances across the views **G**, while the minimal distance is not present in **G**_∧_. In that case we randomly pick one view. Consequently, the algorithm needs to run multiple times in order to deliver an adequate estimate of the view-specific contributions to the data fusion operation. We introduce the parameter `HC.iter` which is set to a minimum limit of 10 as a default.

#### Algorithm 1: Data fusion with hierarchical clustering

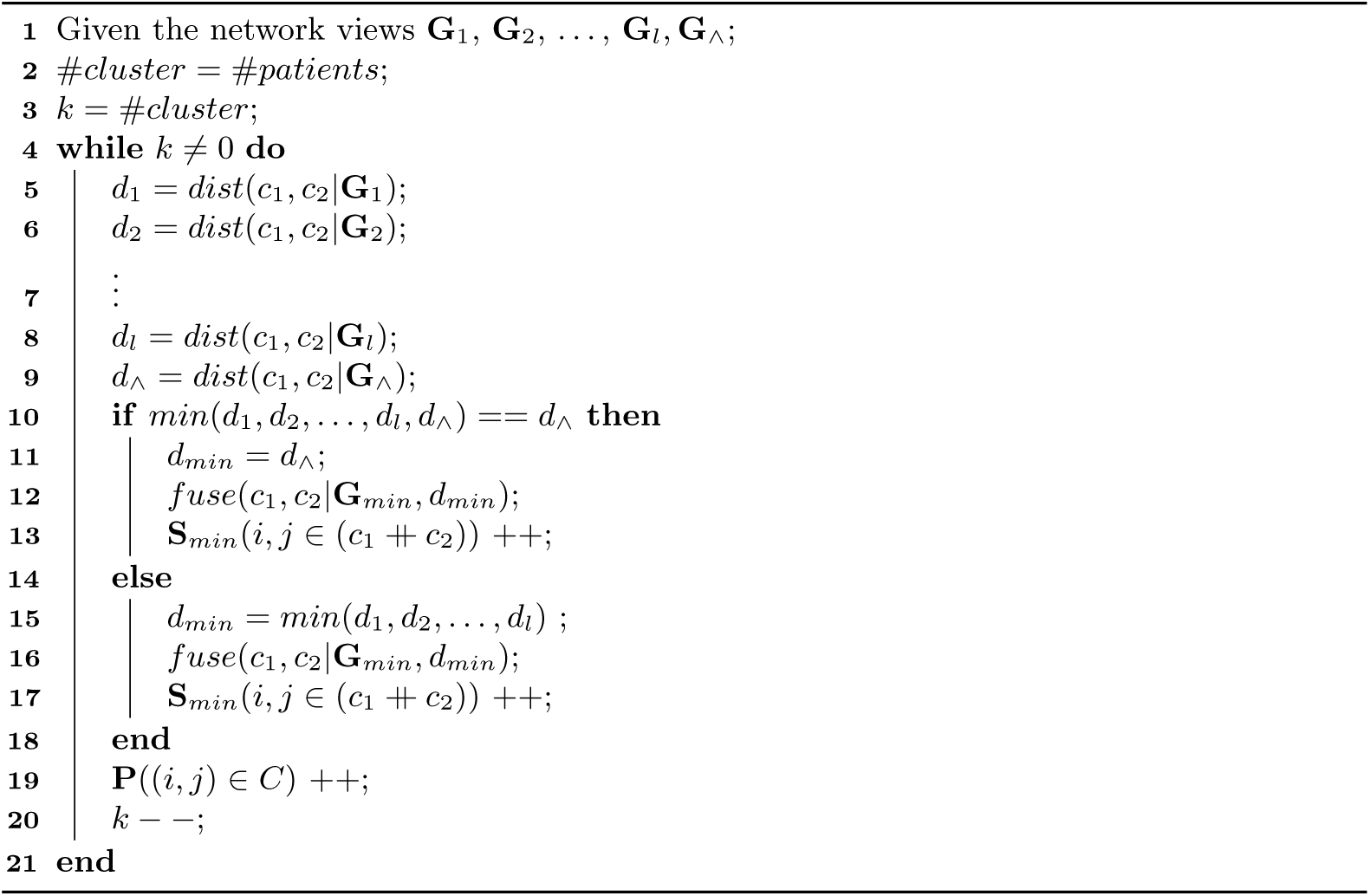

### 2.5 Simulation 1: Disjoint inter-cluster elements

In a first simulation we generate two data views represented as numerical matrices (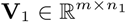 and 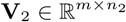). The first matrix reflects three clusters, sampled from Gaussian distributions, *c*_1_ = 𝒩 (−10, *σ*^2^), *c*_2_ = 𝒩 (0, *σ*^2^), and *c*_3_ = 𝒩 (10, *σ*^2^). Each of the three clusters contains four elements. The numbers of features are *n*_1_ = 100 in support of these cluster assignments. For the second data view we generate two cluster, *c*_1_ = 𝒩 (0, *σ*^2^) and *c*_2_ = 𝒩 (10, *σ*^2^). In this case the number of features is *n*_2_ = 1000. The two views are denoted as follows:

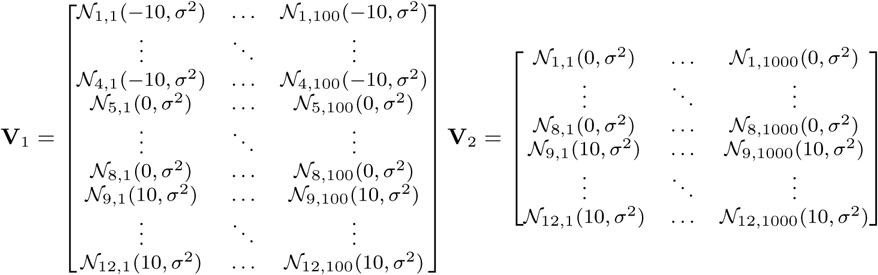

It is evident from these views that the first two cluster in **V**_1_ ({1, …, 4} and {5, …, 8}) form a subset of the first cluster in **V**_2_ ({1, …, 8}). After data integration we expect a final cluster solution of three clusters (*c*_1_ = {1, …, 4}, *c*_2_ = {5, …, 8}, and *c*_1_ = {9, …, 12}) because these clusters are supported by both views. However, since cluster *c*_1_ and *c*_2_ are fully connected in the second view, we expect these two clusters to be closer to each other than to *c*_3_.

We vary the parameter *σ*^2^ = [0.1, 0.5, 1, 5, 10, 20] and expect that the cluster quality decreases the higher the variances for the specific groups. The effect of *σ*^2^ on the clusters is graphical illustrated in Supplementary Figure 1. We analyze how *HC-fused* is affected by these variations accordingly.

### 2.6 Simulation 2: Disjunct inter-cluster elements

For the second simulation we formulate two views, 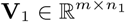 and 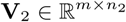. The first view reflects three clusters, *c*_1_ = 𝒩 (−10, *σ*^2^), *c*_2_ = 𝒩 (0, *σ*^2^), and *c*_3_ = 𝒩 (10, *σ*^2^). In this case, the first and the third cluster contains two elements, respectively, and the second cluster six elements. The second view represents two clusters, *c*_1_ = 𝒩 (−10, *σ*^2^) and *c*_2_ = 𝒩 (0, *σ*^2^), plus two single elements, *c*_3_ = 𝒩 (10, *σ*^2^) and *c*_4_ = 𝒩 (30, *σ*^2^). The only difference between **V**_1_ and **V**_2_ is that in view **V**_2_ the last two elements do not form a cluster.

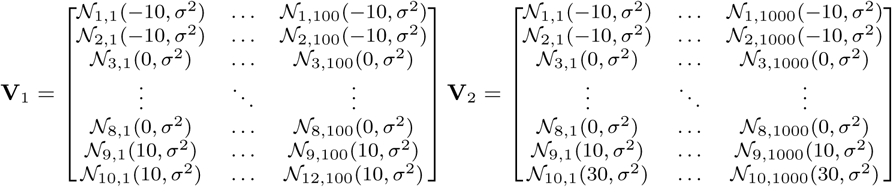

After data integration we expect a final solution of three clusters (*c*_1_ = {1, 2}, *c*_2_ = {3, …, 8} and *c*_3_ = {9, 10}). The clusters *c*_1_ and *c*_2_ should be inferred with high confidence because these are confirmed in both data views. A lower confidence should be assigned to the third cluster *c*_3_ because it is just confirmed in the first view. Again, we vary the parameter *σ*^2^ = [0.1, 0.5, 1, 5, 10, 20]. The effect of *σ*^2^ on the clusters is graphical illustrated in Supplementary Figure 2.

### 2.7 Comparison with other methods

We compare *HC-fused* to the *state-of-the-art methods SNF* [8], *PINSPlus* [18], *and NEMO* [9]. In addition, we match the performance of these methods to a baseline approach *(HC-concatenate*) where data are simply concatenated, and a single-omics hierarchical clustering approach based on Ward’s method is applied. We administer the Adjusted Rand Index (ARI) [21] as a performance measure and the *Silhouette Coefficient* (SIL) [22] for cluster quality assessment. Figures are generated using the R-package *ggnet* [23] and *ggplot2* [24]. Simulation results can be easily reproduced by the R-scripts provided in our GitHub repository (pievos101/HC-fused).

## 3 Results

### 3.1 Disjoint inter-cluster elements

The results from the simulations with disjoint inter-cluster elements are illustrated in Fig. 1. *HC-fused* infers three clusters for the first view and two clusters for the second view. At this stage, the similarity weightings of the vertices are all equal to 1. After applying our proposed algorithm for data fusion, a fused network is constructed as seen in Fig. 1C. The optimal number of clusters is *k* = 3, as inferred by the *Silhouette Coefficient* based on the matrix **P**. Panel D shows the dendrogram when the hierarchical clustering algorithm based on Ward’s method is applied to the fused matrix **P**. The cluster elements {9, …, 12} are most distant to the other elements because they are disconnected from these elements in both views. The elements within the three clusters are all equally distant to each other because all connections within these clusters are confirmed in the **G**_∧_ view.

**Fig 1.**
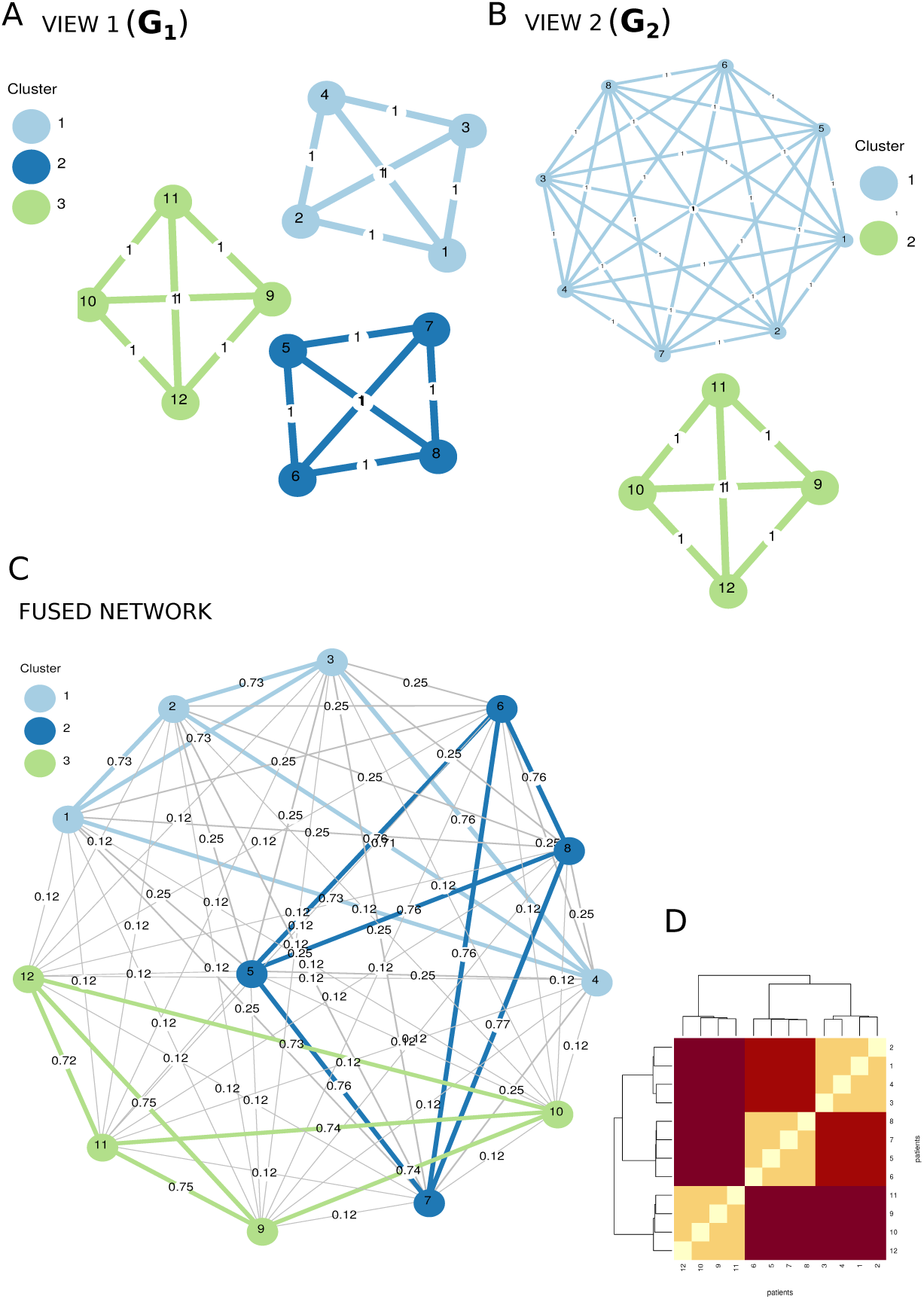
Results from simulation 1 (disjoint inter-cluster elements with *σ*^2^ = 1). **A**. Shown is **G**_1_ as a result from the first view (**V**_1_). **B**. Shown is **G**_2_ as a cluster result from the second view (**V**_2_). **C**. The fused network based on the fused similarity matrix **P**. Three clusters are suggested by the *Silhouette Coefficient*. **D**. The resulting dendrogram when hierarchical clustering is applied to the fused similarity matrix **P**.

Fig. 2 highlights the contributions of the views to the data fusion process. The cluster members {9, …, 12} are fully supported by the **G**_∧_ view, whereas the view **G**_2_, also contributes to the other elements. This is not surprising because the concerned elements are all connected in the second view.

**Fig 2.**
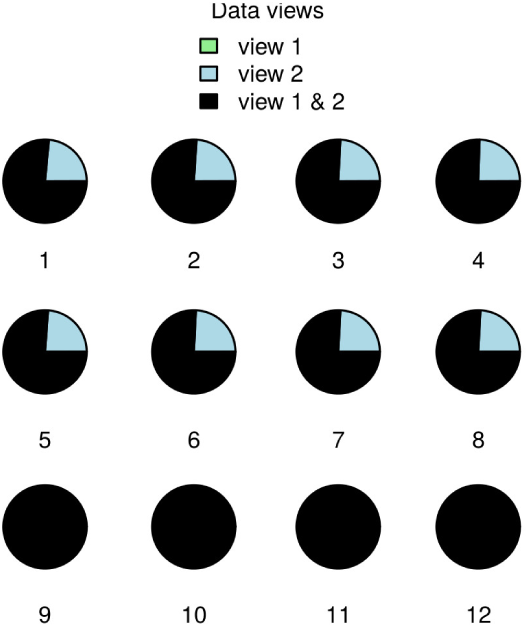
Results for simulation 1. Contribution of the views to the hierarchical data fusion.

It can be seen that *HC-fused* is competing well with the *state-of-the-art* methods (Fig. 3). To our surprise, *SNF* performs very weak. The *eigen-gaps* method, as mentioned by its authors, infers two cluster as the optimal solution. It does not take into account that cluster *c*_1_ and cluster *c*_2_ are disconnected in the first view. Also, the *Silhouette* method applied to the fused affinity matrix infers only two cluster. We observe similar ARI values for *HC-fused, PINSPlus*, and *NEMO*. Compared to *HC-fused, PINSPlus* and *NEMO* are more robust against increased within-cluster variances.

**Fig 3.**
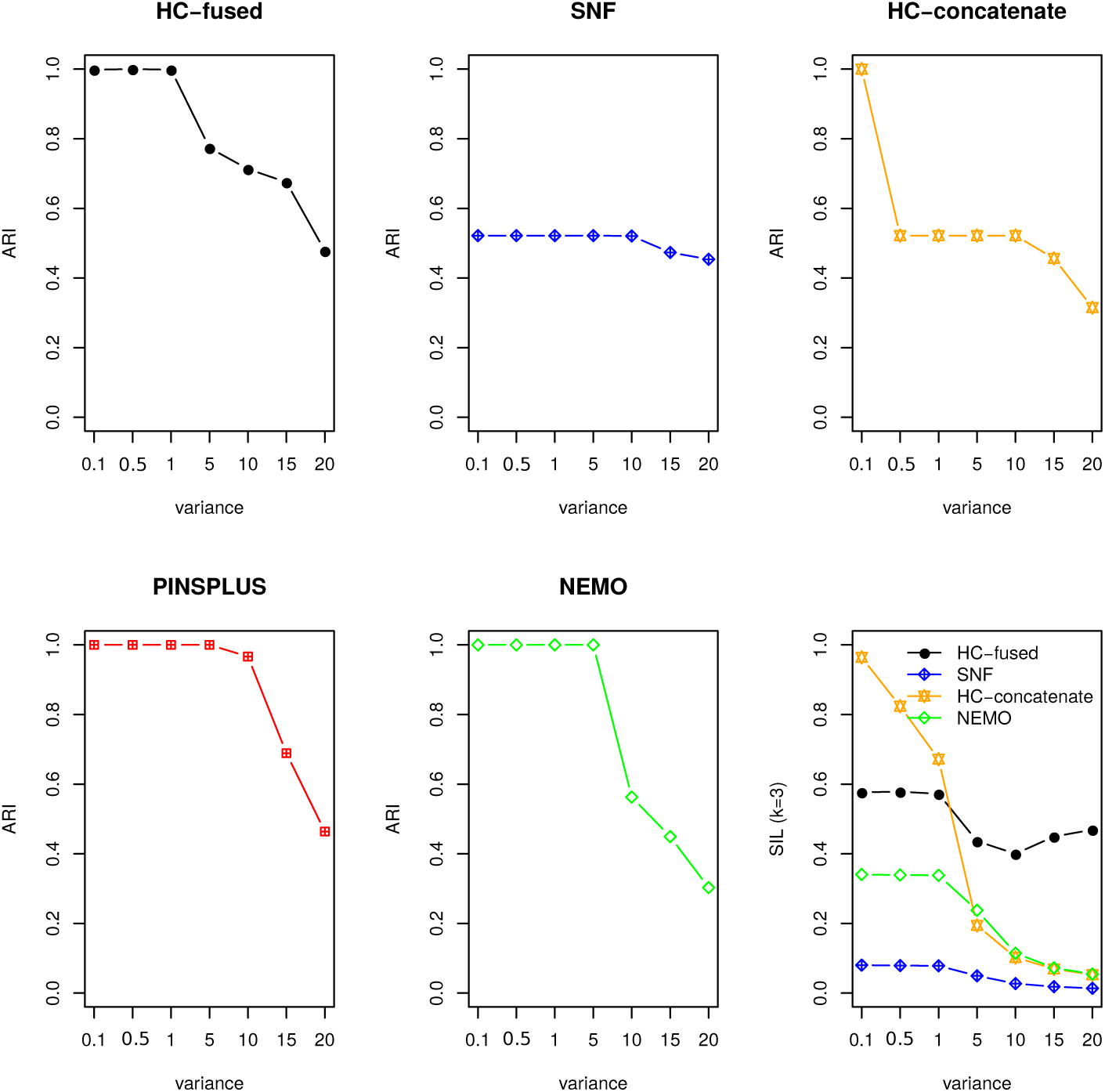
Results from simulation 1 (disjoint inter-cluster elements with *σ*^2^ = [0.1, 0.5, 1, 5, 10, 15, 20]). We compare *HC-fused* with *SNF, PINSPlus, NEMO*, and *HC-concatenate*. The true number of clusters is *k* = 3, with the cluster assignments *c*_1_ = {1, …, 4}, *c*_2_ = {4, …, 8}, and *c*_3_ = {9, …, 12}. The performance is measured by ARI. For each *σ*^2^ 100 runs were performed and the mean ARI values are displayed. The panel in the bottom right shows the mean SIL coefficients for the true cluster assignments (*k* = 3).

Starting with a within-cluster variance of *σ*^2^ = 1, *HC-fused* frequently behaves like *SNF* and infers the cluster assignments as represented by the second view (Fig. 1). Simply concatenating the data views (*HC-concatenate*) has the consequence of an overall low accuracy. This is expected because the second view contains 10 times more features and thus gives more weights to the cluster assignments in **V**_2_. The SIL coefficient, as a measure of cluster quality, is highest for the *HC-concatenate* approach for low to medium variances. However, it suggests even higher SIL values for a *k* = 2 cluster solution. The SIL coefficient of *HC-fused* is markedly higher than those of *SNF* and *NEMO*. We cannot give any cluster quality measure for *PINSPlus* because the corresponding R-package does not provide a single fused distance or a similarity matrix.

### 3.2 Disjunct inter-cluster elements

The hierarchical fusion process via *HC-fused* is illustrated in Fig. 4. The only difference between the two network views shown in Fig. 4A and Fig. 4B is, that in Fig. 4B the elements 9 and 10 are not connected. After data fusion (Fig. 4C) the SIL coefficient infers three clusters as the optimal solution. The cluster elements in *c*_3_ = {9, 10} are substantially distant from each other (Fig. 4D) and signify a contribution from view 1, as shown in Fig. 5. This is expected because they are only connected in the first view and thus the confidence about the relevance of this cluster is reduced. The cluster elements *c*_2_ = {3, …, 8} are mainly fused in matrix *G*_∧_ because the cluster is confirmed by both views (Fig. 5). The same applies to the cluster *c*_1_ = {1, 2} and thus the elements within *c*_1_ and *c*_2_ have equal distances to each other (Fig. 4C, 4D).

**Fig 4.**
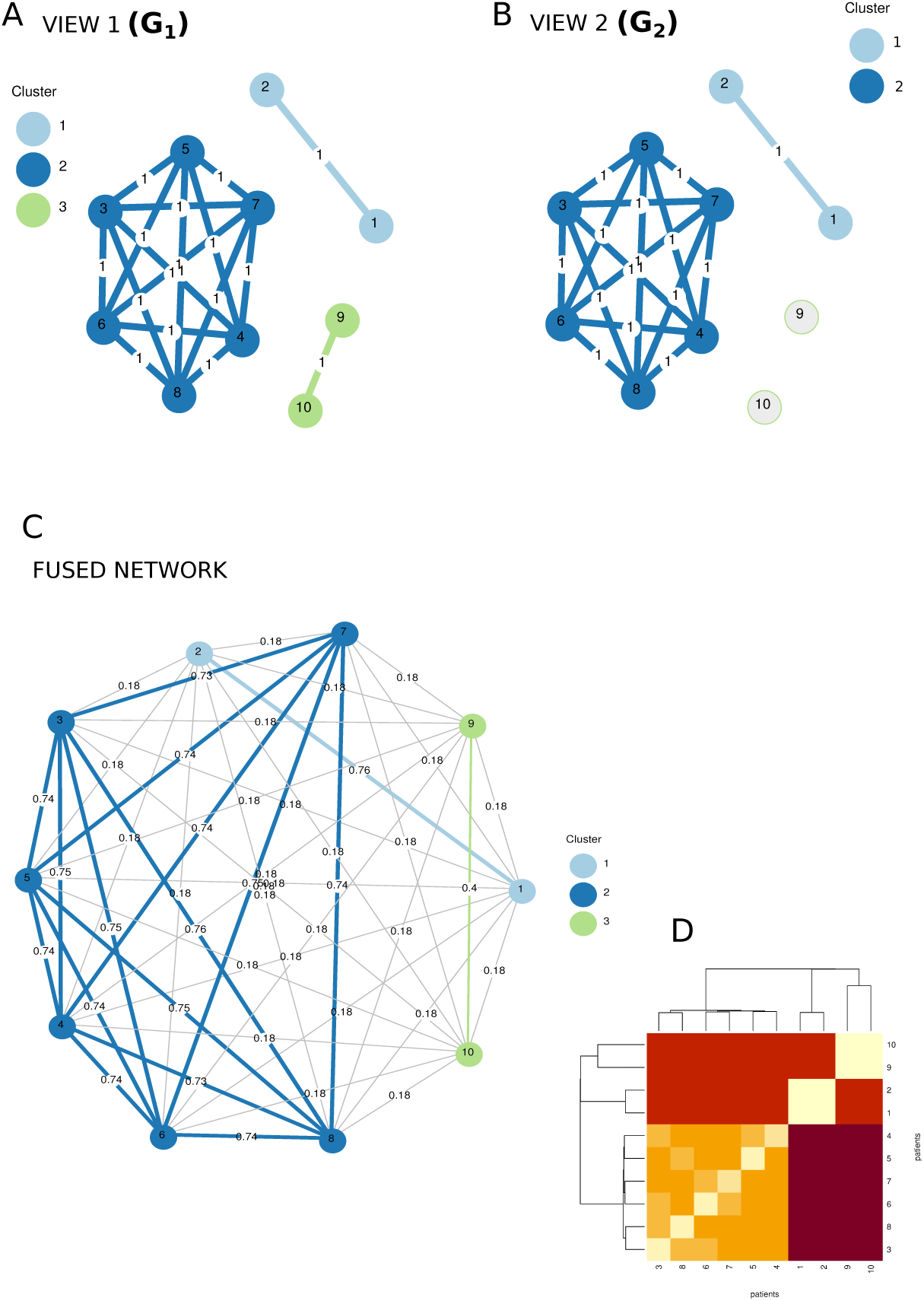
Results from simulation 2 (disjunct inter-cluster elements with *σ*^2^ = 1). **A**. Shown is **G**_1_ as a cluster result from the first view (**V**_1_). **B**. Shown is **G**_2_ as a cluster result from the second view (**V**_2_). **C**. The fused network based on the fused similarity matrix **P**. Three clusters are suggested by the SIL coefficient. **D**. The resulting dendrogram when hierarchical clustering is applied to the fused similarity matrix **P**.

**Fig 5.**
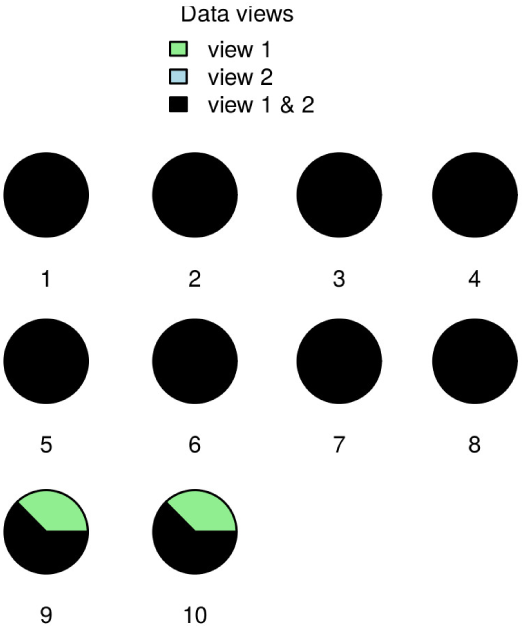
Results for simulation 2. Contribution of the views to the hierarchical data fusion.

Overall, the results of the simulation with disjunct cluster elements are the best for *HC-fused. PINSplus* cannot compete with *HC-fused* (Fig. 4), it constantly infers four clusters as the optimal solution and does not take into account the connectivity between elements 9 and 10 in the first view. Starting with a within-cluster variance of *σ*^2^ = 1, the same can be observed for *HC-fused* (see Fig. 4). *NEMO* performs surprisingly weak. The modified *eigen-gap* method, as pointed out by the authors, performs poorly in this specific simulation scenario. *NEMO* infers far more than three clusters and the elements seem to be randomly connected to each other. When reducing the number of neighborhood points in the diffusion process, the total number of clusters is slightly decreasing but with no relevant gain in accuracy. Interestingly, with the same number of neighborhood points *SNF* performs much better. In a further investigation, when the *Silhouette* method is adopted to the fused similarity matrix from *NEMO*, the true number of clusters can be obtained. This fact points at a potential problem with the *eigen-gap* method as implemented in *NEMO* for data sets with disjunct inter-cluster elements.

When conducting cluster quality assessments, again we can observe low SIL values for the fused affinity matrix resulting from *SNF* (see Fig. 6). For low to medium within-cluster variances, results by *NEMO* are comparable to those by *HC-fused*. A value for cluster quality cannot be reported for *PINSPlus* because no single fused data view is available.

**Fig 6.**
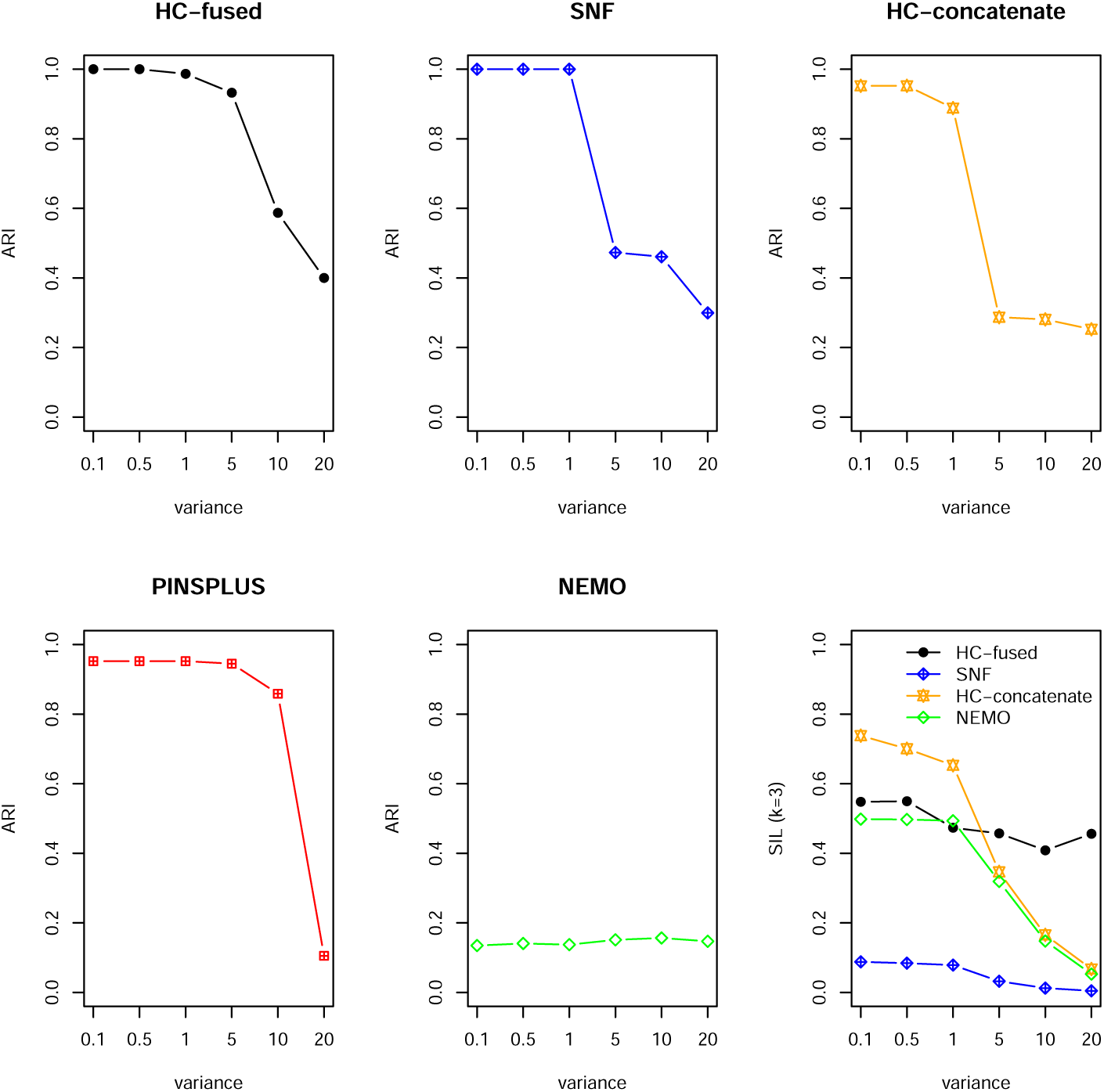
Results from simulation 2 (disjunct inter-cluster elements with *σ*^2^ = [0.1, 0.5, 1, 5, 10, 15, 20]). We compare *HC-fused* with *SNF, PINSPlus, NEMO* and *HC-concatenate*. The true number of clusters is *k* = 3, with the cluster assignments *c*_1_ = {1, 2}, *c*_2_ ={3, …, 8}, and *c*_3_ = {9, 10}. The performance is measured by ARI. For each *σ*^2^ 100 runs were performed and the mean ARI values are shown. The panel at the bottom right displays the mean SIL coefficients for the true cluster assignments (*k* = 3).

### 3.3 Disjoint & disjunct inter-cluster elements

In addition to the above studied simulation scenarios, which represent two fundamentally different cluster patterns across the views, we also studied a mixture of both. We simulated two views comprising disjoint and disjunct inter-cluster elements. This particular simulation scenario is described in detail in the supplementary material. We observed that *HC-fused* is clearly outperforming the competing methods (Supplementary Fig. 5). To our surprise, none of the *state-of-the-art* methods (*SNF, NEMO*, and *PINSPlus*) infers the correct number of clusters, even when the *within-cluster* variances are rather low. The results obtained from *PINSPlus* are almost as good as from *HC-fused*. For medium-size variances *PINSPlus* does slightly better. The cluster quality of *HC-fused*, as expressed by the SIL coefficients, is higher than those of *NEMO* and *SNF* (Supplementary Fig. 5, bottom right).

### 3.4 Robustness analysis

Sets of features, in size 1,5,10, and 50, were randomly permuted across the objects in order to test the stability of the approaches to predict the correct cluster solution. One hundred runs were executed for each setting and the mean ARI values calculated. As can be seen from the Supplementary Fig. 6, *NEMO* and *PINSPlus* are more stable against noise compared to *HC-fused*. When the number of permuted features is greater than one, the accuracy of *HC-fused* drops. This is most likely due to the fact that *HC-fused* uses the Euclidean distance to generate the connectivity matrices **G**. It is well known that the Euclidean distance is prone to outliers. Removing concerned data points prior to the analysis may be a necessary initial step. Another possible approach would be the application of principal component analysis (PCA) on the feature space.

In case of disjunct cluster elements (Supplementary Fig. 7) we observe a slightly different outcome situation. *HC-fused* is definitely more robust against noise compared to *SNF. PINSPlus* provides also stable results, but as already pointed out in the previous section, produces a wrong cluster assignment.

### 3.5 Application of integrative clustering to TCGA cancer data

To demonstrate the usability of our approach we applied it to the TCGA cancer data as provided by [16]. TCGA provides mRNA, methylation, and miRNA data for a fixed set of patients. We tested our approach *HC-fused* on nine different cancer types: glioblastoma multiforme (GBM), kidney renal clear cell carcinoma (KIRC), colon adenocarcinoma (COAD), liver hepatocellular carcinoma (LIHC), skin cutaneous melanoma (SKCM), ovarian serous cystadenocarcinoma (OV), sarcoma (SARC), acute myeloid leukemia (AML), and breast cancer (BIC).

In contrast to other benchmark studies, that apply multi-omics approaches to a static data set, we randomly sample 20 times 100 patients from the data pool, performed survival analysis and calculated the Cox log-rank [25]. The thus obtained *p*-values are summarized in boxplots in Fig. 7. We are convinced that our approach is conveying a less biased picture of the clustering performance. In summary, we observe an overall weaker performance across all methods than previously reported in [16] (Fig. 7, Supplementary Table 1). *HC-fused* performs best for KIRC, LIHC, SKCM, OV, and SARC when the median log-rank *p*-values are used for comparison. Global best results are observed for the KIRC and SARC data sets. The method implemented in the R-package *NEMO* outperforms the other methods for the GBM and AML cancer types. *PINSplus* shows an overall low performance in almost all cases. Notably, all methods studied here perform weak on the COAD data set. Above observations indicate substantial differences in structure and quality of the various cancer type data.

**Fig 7.**
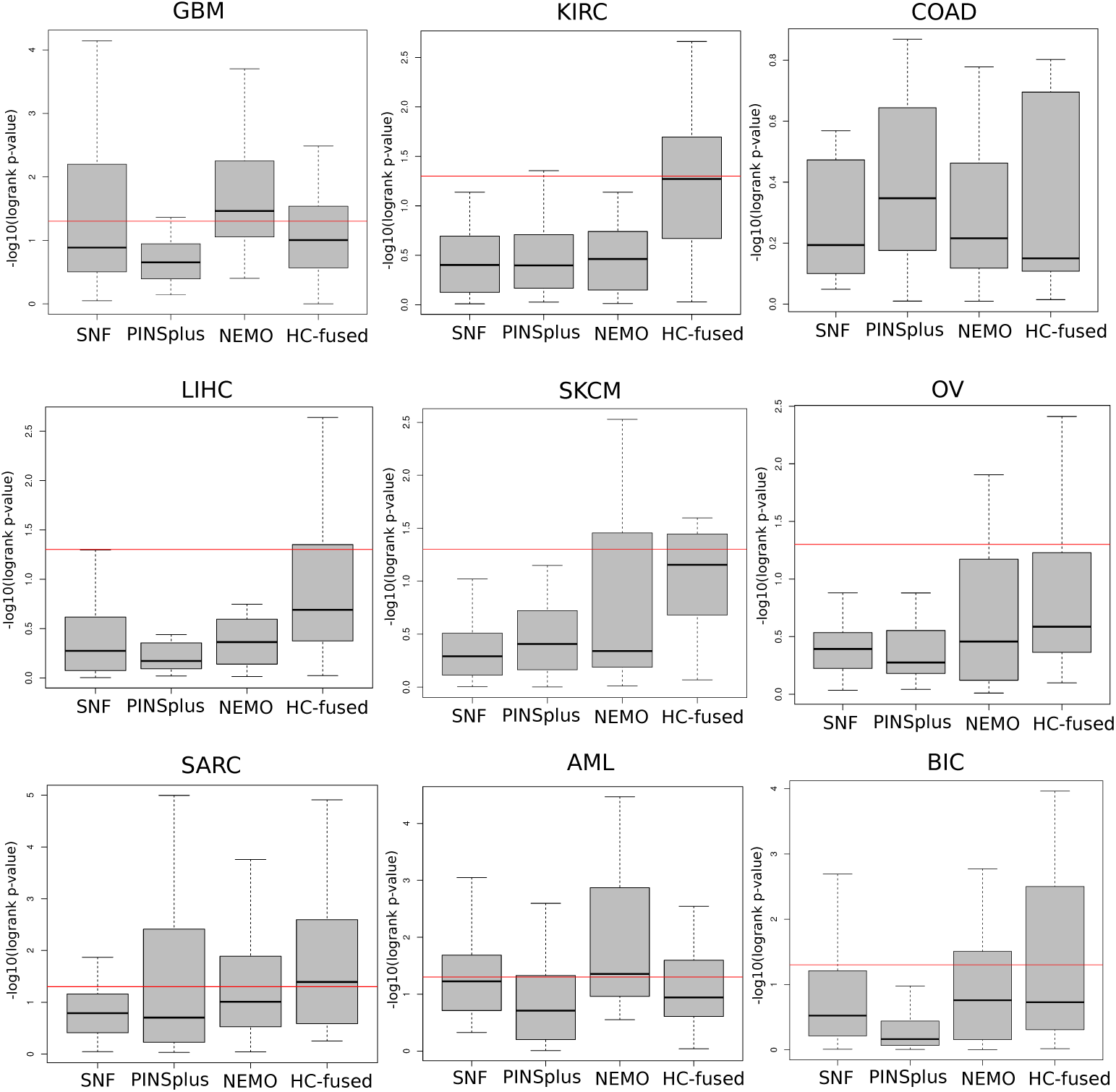
TCGA integrative clustering results. Displayed are the log-rank *p*-values on a logarithmic scale for nine different cancer types. The red line refers to *α* = 0.05 significance level.

Based on the analysis that produced the findings in Fig. 7, we applied our approach to the TCGA breast cancer data as allocated by [8]. In the supplementary material we provide a *step-by-step* guide on how to analyze this data set within the R environment using *HC-fused. HC-fused* infers seven clusters (Supplementary Fig. 8) as the optimal solution with a significant Cox log-rank *p*-value of 3.2^−5^. Previously reported *p*-values for the same data set take the values *p* = 1.1^−3^ for *SNF* and *p* = 3.0^−4^ for *rMKL-LPP*. Clusters 1,2,4, and 5 are mainly confirmed by all biological layers, whereas cluster 3,6, and 7 signify some exclusive contributions from the single-omics views to the data fusion process. Especially the mRNA expression data seem to substantially contribute to clusters 3 and 7 (Supplementary Fig. 9).

## 4 Discussion

In this article, we have developed a hierarchical clustering approach for multi-omics data fusion based on simple and well established concepts. Simulations on *disjoint* and *disjunct* cluster elements across simulated data views indicate superior results over recently published methods. In fact, we provide two simulation scenarios in which *state-of-the-art* methods behave remarkably different. We discovered that *NEMO* performs well on data sets with *disjoint inter-cluster elements*, whereas *SNF* does much better on *disjunct inter-cluster elements* across the data views. We hope that our synthetic data sets may act as a useful benchmark for future studies in the field.

The application on real multi-omics TCGA cancer data suggest promising results of *HC-fused*. It competes well with *state-of-the-art* methods. Out of nine studied cancer types, HC-fused performs best in five cases. Importantly and in contrast to other approaches, *HC-fused* provides information about the contribution of single-omics data to the data fusion process. It should be noted, however, that our algorithm requires multiple iterations to achieve a high-quality estimate. The reason is that in each integration step the single views may contain comparable minimal distances. As a consequence, there might be uncertainty about the views for which data points should be fused. Currently, we solve this problem by a uniform sampling scheme plus running the proposed algorithm multiple times. At the moment it is not entirely clear how many iteration steps should be used. We suggest to apply as minimum `HC.iter`>= 10. Values above this minimum have produced reasonable results in our investigations. We plan to solve this problem in a computational more feasible way in the next release of the *HC-fused* R-package. A promising approach would be to model the fusion algorithm as a Markov process where each view represents a state and the transition probabilities depend on the number of view-specific items providing the same minimal distance.

Unlike other approaches, the *HC-fused* workflow does not depend on a specific clustering algorithm. This means, with the current release, any hierarchical clustering method provided by the native R function hclust can be used to create the connectivity matrices **G**. Also, the final fused matrix *P* can be calculated by an arbitrary clustering algorithm preferred by a user. Further investigations are needed to study popular cluster algorithms within the proposed

*HC-fused* workflow to learn how they might influence the fusion results. Given the known heterogeneity of omics data, this might be worthwhile to work on next. Another characteristic of *HC-fused* is its independence of a specific technique to infer the best number of clusters. While, in this work, the *Silhouette Coefficient* was adopted, other assessment parameters might improve the outcomes. We plan to further develop the corresponding GitHub R-package (pievos101/HC-fused), add functionality, and at the same time make sure that the package remains flexible and versatile.

## 5 Conclusion

In this article we have proposed a novel hierarchical data fusion approach embedded in the versatile R-package *HC-fused* available on GitHub (pievos101/HC-fused). Simulations and an application to real-world TCGA cancer data indicate that *HC-fused* is more accurate than or at least as accurate as the *state-of-the-art* methods. In contrast to other approaches, it naturally reports on the contribution of the single-omics data to the data fusion process. Its overall conceptual simplicity fosters the interpretability of the final results. With respect to the biomedical application, multi-omics clustering approaches like the one introduced in this article have certainly the potential to improve the discovery of cancer subtypes. In the future they are most likely to facilitate the treatment of cancer patients in personalized routines.

## Supporting information

HC-fused Supplement

## Acknowledgments

We are grateful to the Kurt und Senta Herrmann-Stiftung, Vaduz, Liechtenstein, for its support. We also thank Luca Vitale and José Antonio Vera-Ramos for helpful discussions.

