## Supplementary material for "A hierarchical clustering and data fusion approach for disease subtype discovery": HC-fused Supplement

May 20, 2020

### 1 Simulation 1: Disjoint inter-cluster elements

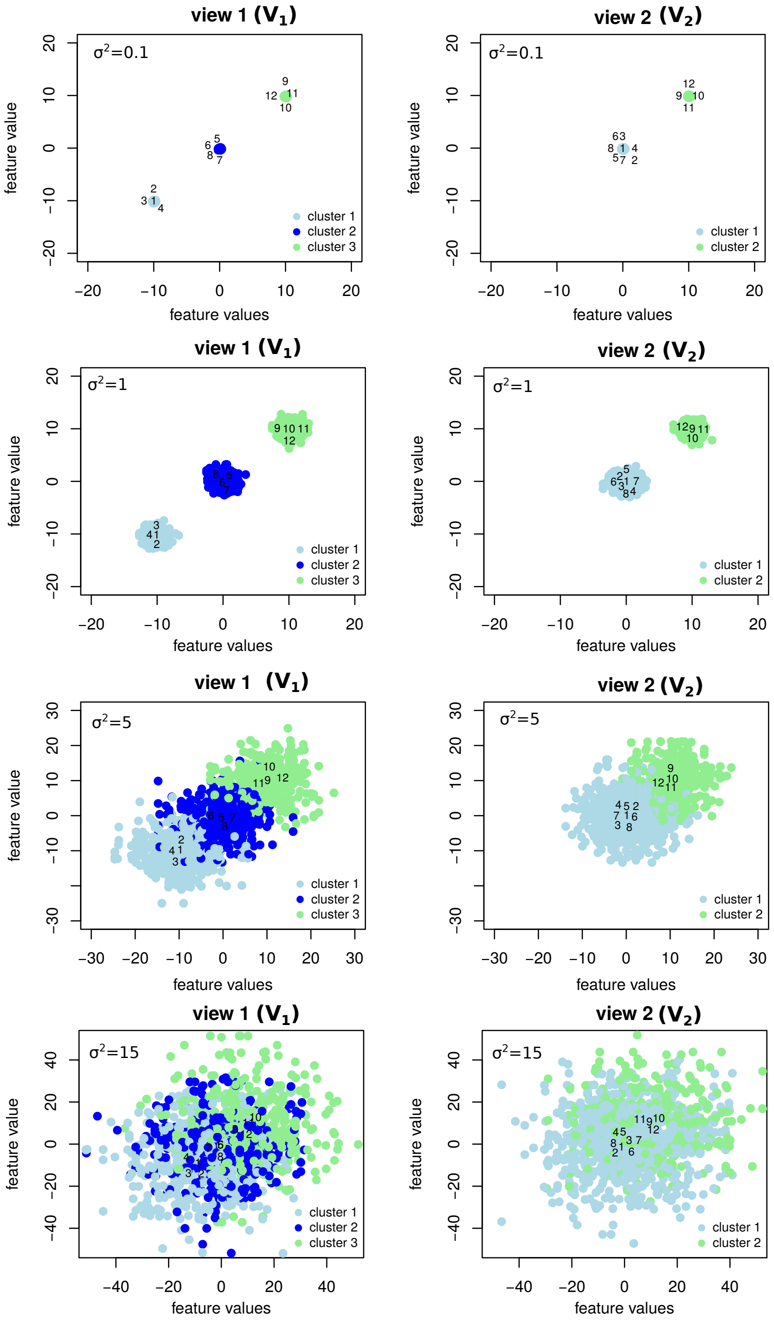


**Supplementary Fig. 1: Disjount inter-cluster elements. The effect of $\sigma^2$ on the view-specific clusters.**

#### 2 Simulation 2: Disjunct inter-cluster elements

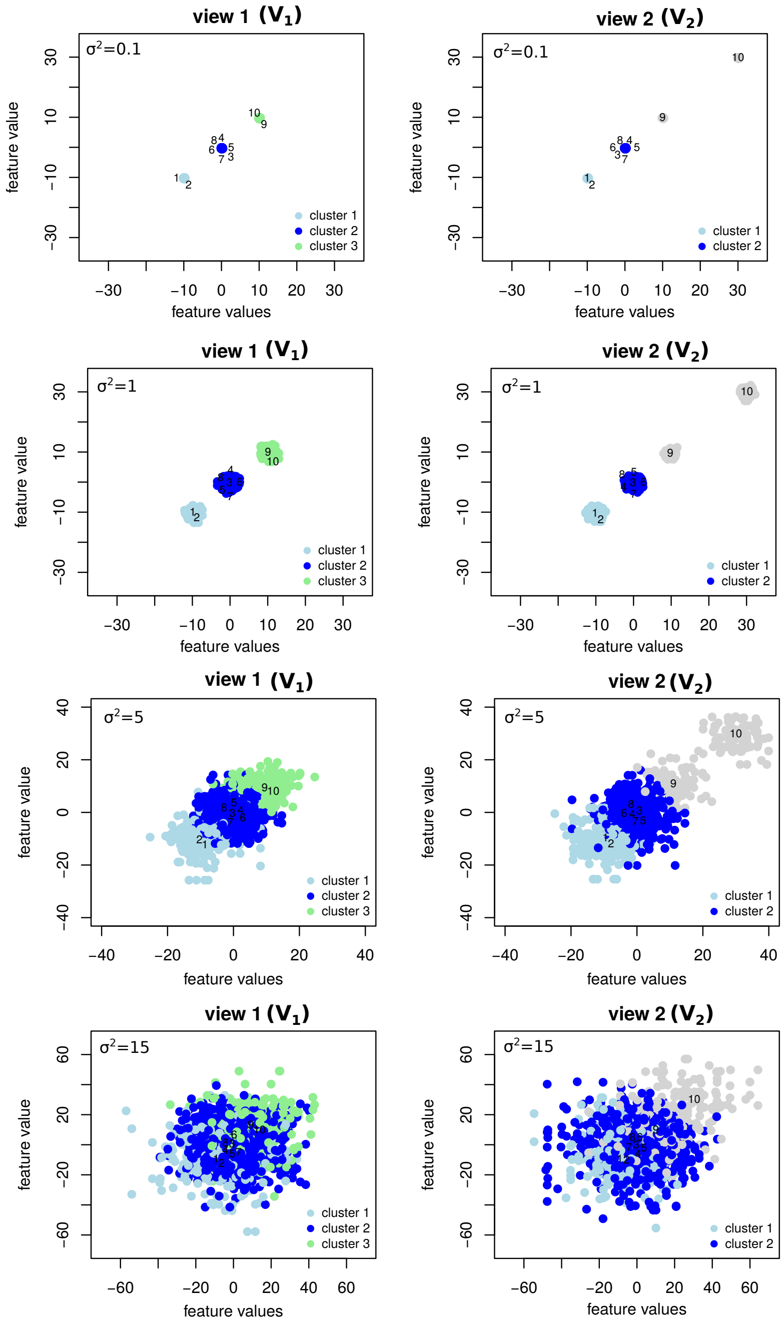

**Supplementary Fig. 2: Disjunct inter-cluster elements. The effect of $\sigma^2$ on the view-specific clusters.**

### 3 Simulation 3: disjoint & disjunct inter-cluster elements

In a third simulation we include a mixture of disjoint and disjunct inter-cluster elements. We formulate two views $\mathbf{V}_1 \in R^{m \times n_1}$ and $\mathbf{V}_2 \in R^{m \times n_2}$. The first view reflects two clusters ($c_1 = \mathcal{N}(-10, \sigma^2)$ and $c_2 = \mathcal{N}(0, \sigma^2)$). In this case, the first cluster contains eight elements, and the second cluster two elements. The second view reflects four clusters ($c_1 = \mathcal{N}(-10, \sigma^2)$, $c_2 = \mathcal{N}(0, \sigma^2)$, $c_3 = \mathcal{N}(10, \sigma^2)$ and $c_4 = \mathcal{N}(30, \sigma^2)$). The difference between $\mathbf{V}_1$ and $\mathbf{V}_2$ is that in view $\mathbf{V}_2$ the last two elements are not connected and the first cluster $c_1 = \{1, 2\}$ is disconnected from all other elements.

$$\mathbf{V}_1 = \begin{bmatrix} \mathcal{N}_{1,1}(-10,\sigma^2) & \dots & \mathcal{N}_{1,1000}(-10,\sigma^2) \\ \vdots & \ddots & \vdots \\ \mathcal{N}_{8,1}(-10,\sigma^2) & \dots & \mathcal{N}_{8,1000}(-10,\sigma^2) \\ \mathcal{N}_{9,1}(0,\sigma^2) & \dots & \mathcal{N}_{9,1000}(0,\sigma^2) \\ \mathcal{N}_{10,1}(0,\sigma^2) & \dots & \mathcal{N}_{12,1000}(0,\sigma^2) \end{bmatrix} \mathbf{V}_2 = \begin{bmatrix} \mathcal{N}_{1,1}(-10,\sigma^2) & \dots & \mathcal{N}_{1,100}(-10,\sigma^2) \\ \mathcal{N}_{2,1}(-10,\sigma^2) & \dots & \mathcal{N}_{2,100}(-10,\sigma^2) \\ \mathcal{N}_{3,1}(0,\sigma^2) & \dots & \mathcal{N}_{3,100}(0,\sigma^2) \\ \vdots & \ddots & \vdots \\ \mathcal{N}_{8,1}(0,\sigma^2) & \dots & \mathcal{N}_{8,100}(0,\sigma^2) \\ \mathcal{N}_{9,1}(10,\sigma^2) & \dots & \mathcal{N}_{9,100}(10,\sigma^2) \\ \mathcal{N}_{10,1}(30,\sigma^2) & \dots & \mathcal{N}_{10,100}(30,\sigma^2) \end{bmatrix}$$

We vary the parameter $\sigma^2 = [0.1, 0.5, 1, 5, 10, 20]$.

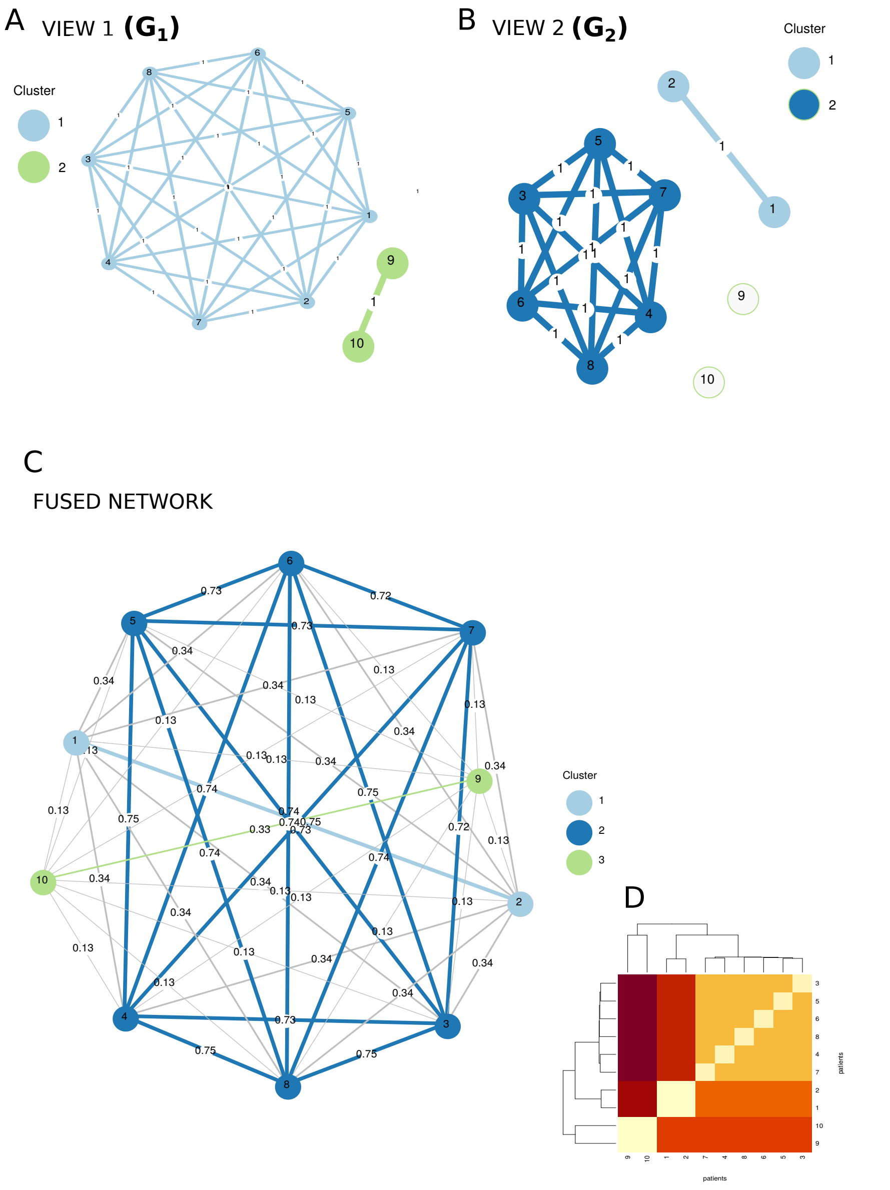


**Supplementary Fig. 3:** Disjount & disjunct inter-cluster elements. **A**. Shown is $\mathbf{G}_1$ from the first view ($\mathbf{V}_1$). **B**. Shown is $\mathbf{G}_2$ from the second view ($\mathbf{V}_2$). **C**. The fused network based on the fused similarity matrix $\mathbf{P}$. Three clusters are suggested by the silhouette coefficient. **D**. The resulting dendrogram when hierarchical clustering based on Ward's method is applied to the fused similarity matrix $\mathbf{P}$.

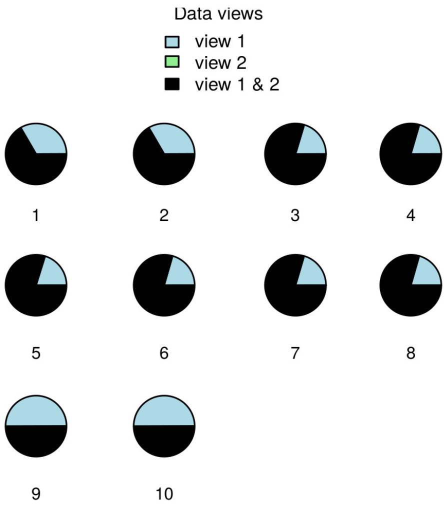


**Supplementary Fig. 4:** Results for disjoint & disjunct inter-cluster elements. Contribution of the views to the hierarchical data fusion.

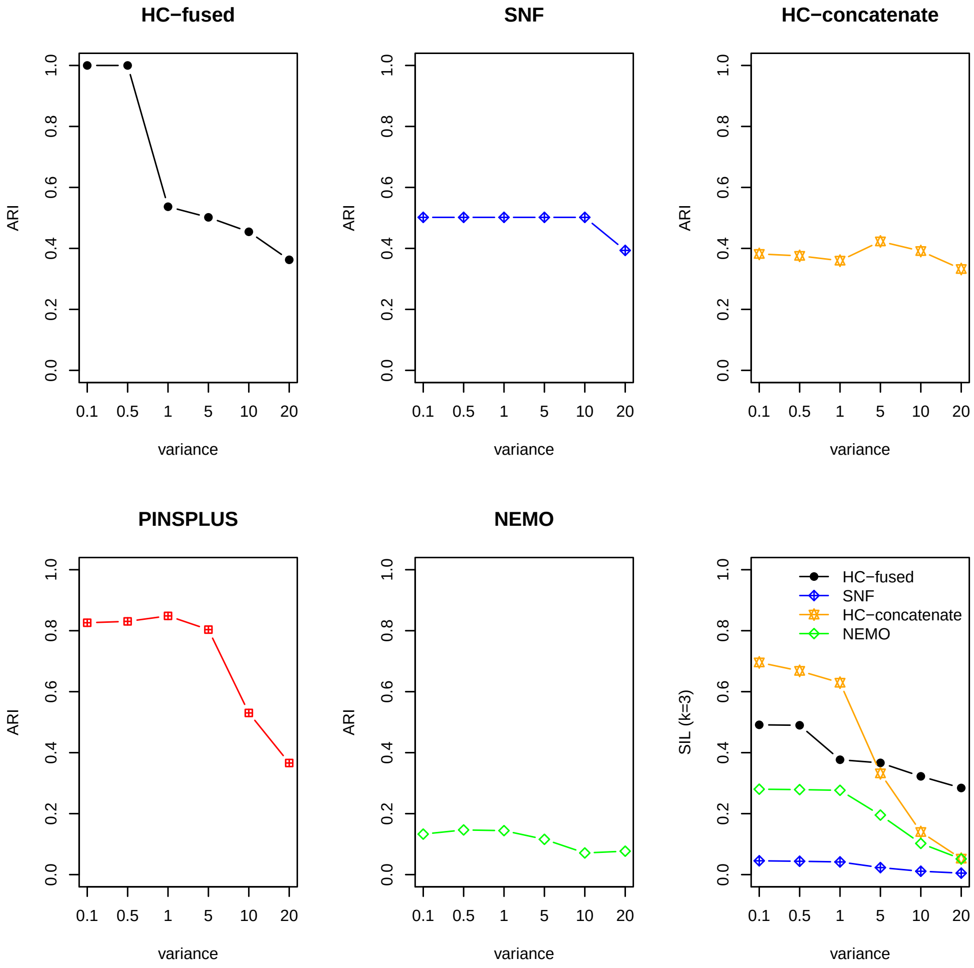


**Supplementary Fig. 5:** Results from simulation 3 (disjunct & disjoint inter-cluster elements with $\sigma^2 = [0.1, 0.5, 1, 5, 10, 15, 20]$). We compare *HC-fused* with *SNF*, *PINSPlus*, *NEMO*, and *HC-concatenate*. The true number of clusters is $k = 3$, with the cluster assignments $c_1 = \{1, 2\}$, $c_2 = \{3, \ldots, 8\}$, and $c_3 = \{9, 10\}$. For each $\sigma^2$ we performed 100 runs and show the mean ARI value. The panel at the bottom right displays the SIL coefficients for the true cluster assignments ($k = 3$).

### 4 Robustness analyses

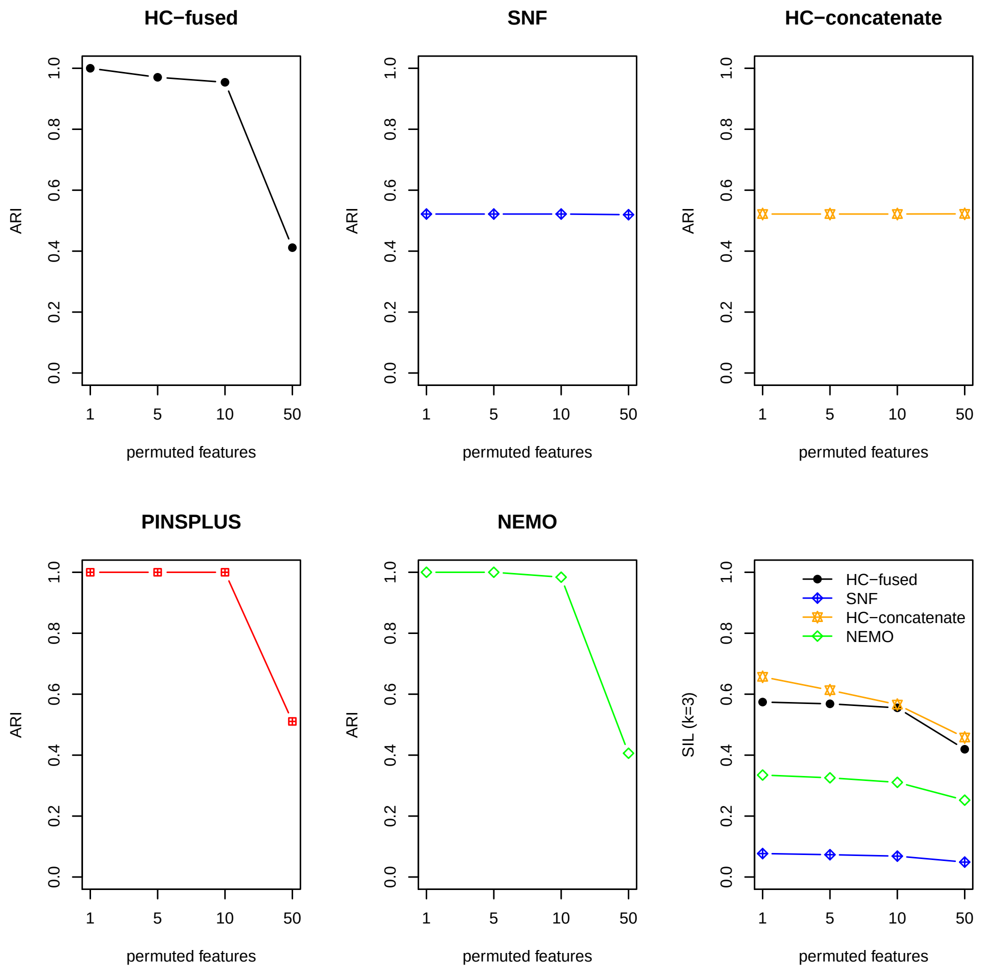


**Supplementary Fig. 6:** Robustness analyses for simulation 1 (disjoint inter-cluster elements with $\sigma^2 = 1$). We compare *HC-fused* with *SNF*, *PINSPlus*, *NEMO*, and *HC-concatenate*. The true number of clusters is $k = 3$, with the cluster assignments $c_1 = \{1, 2\}$, $c_2 = \{3, \ldots, 8\}$, and $c_3 = \{9, 10\}$. We randomly permuted a set of features (1, 5, 10 and 50) to test the methods on their robustness to predict the true cluster assignments. We performed 100 runs for each setting and show the mean ARI value. The panel at the bottom right displays the SIL values for the true cluster assignments ($k = 3$).

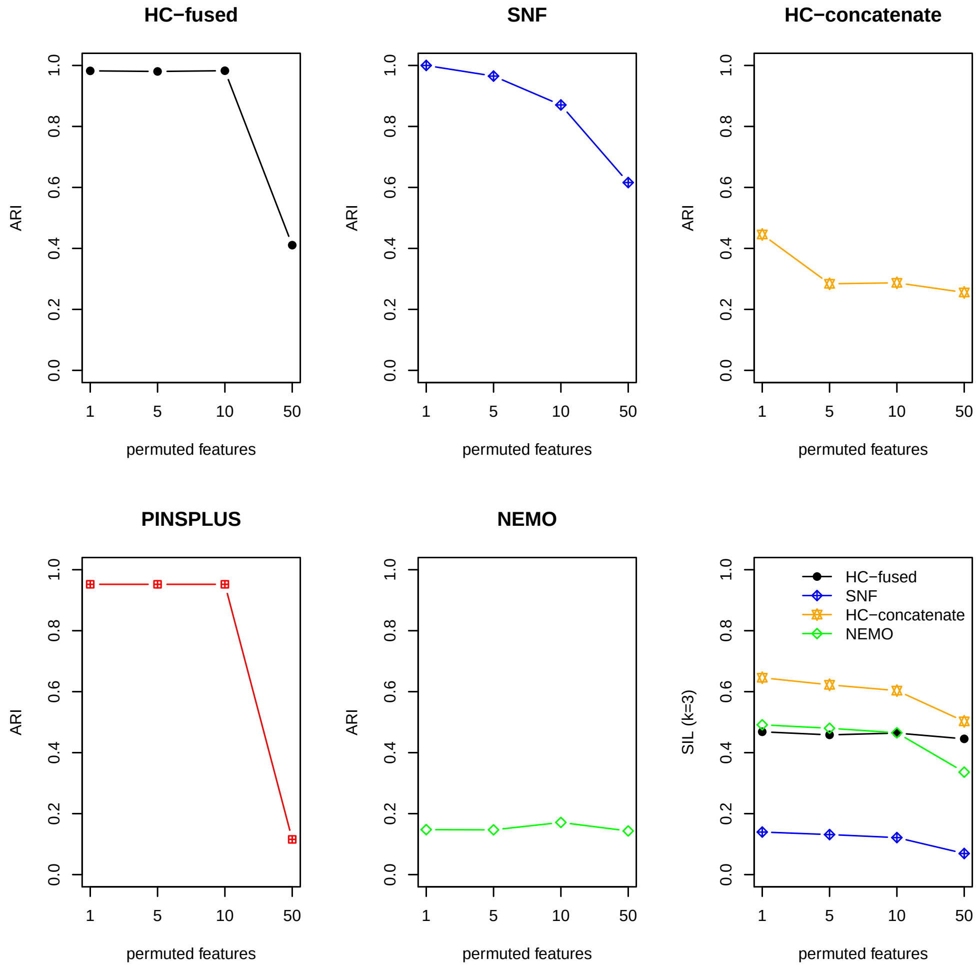


**Supplementary Fig. 7:** Robustness analyses for simulation 2 (disjunct inter-cluster elements with $\sigma^2 = 1$). We compare *HC-fused* with *SNF*, *PINSPlus*, *NEMO*, and *HC-concatenate*. The true number of clusters is $k = 3$, with the cluster assignments $c_1 = \{1, 2\}$, $c_2 = \{3, \ldots, 8\}$, and $c_3 = \{9, 10\}$. We randomly permute a set of features (1, 5, 10 and 50) to test the methods on their robustness to predict the true cluster assignments. We performed 100 runs for each setting and show the mean ARI value. The panel at the bottom right displays the SIL coefficients for the true cluster assignments ($k = 3$).

### 5 Integrative clustering of TCGA cancer data

**Supplementary Table 1:** Survival analyses of TCGA cancer data.

| Cancer type | SNF | PINSPLUS | NEMO | HC-fused |
|---|---|---|---|---|
| GBM | 0.1304 | 0.2223 | **0.0347** | 0.0997 |
| KIRC | 0.3962 | 0.4005 | 0.3464 | **0.0561** |
| COAD | 0.6402 | **0.4500** | 0.6092 | 0.7081 |
| LIHC | 0.5357 | 0.6731 | 0.4354 | **0.2062** |
| SKCM | 0.5153 | 0.3956 | 0.4565 | **0.0699** |
| OV | 0.4042 | 0.5300 | 0.3593 | **0.2594** |
| SARC | 0.1622 | 0.2024 | 0.0979 | **0.0408** |
| AML | 0.0604 | 0.1973 | **0.0440** | 0.1148 |
| BIC | 0.3004 | 0.6836 | **0.1771** | 0.1870 |

We sample 20 times 100 patients from the data pool and display the median of the corresponding Cox log-rank p-values. The best performing method for each cancer type is highlighted in bold.

### 6 The R-package *HC-fused*: an application to breast cancer data

#### 6.1 Install the *HC-fused* R package

```
install.packages("devtools")
library(devtools)
install_github("pievos101/HC-fused")
```

#### 6.2 Loading the libraries

```
library(HCfused)
library(doParallel)
library(foreach)
library(fastcluster)
library(survival)
```

#### 6.3 Read the data

The breast cancer data matrices are available at https://www.nature.com/articles/nmeth.2810 and can be downloaded from the supplementary material.

```
#mRNA
mRNA <- read.table("BREAST_Gene_Expression.txt")
```

```
mRNA <- t(mRNA)
dim(mRNA)
[1]   105 17814

#methylation data
Methy <- read.table("BREAST_Methy_Expression.txt")
Methy <- t(Methy)
rownames(Methy) <- rownames(mRNA)
dim(Methy)
[1]   105 23094

#miRNA
miRNA <- read.table("BREAST_Mirna_Expression.txt")
miRNA <- t(miRNA)
rownames(miRNA) <- rownames(mRNA)
dim(miRNA)
[1] 105 354

#clinical data
clin <- read.table("BREAST_Survival.txt",stringsAsFactors = FALSE,
                header=TRUE)
```

As seen from the calls above, multi-omics cancer data were collected for 105 patients from three biological layers. The mRNA data contains 17814, the methylation data 23094, and the miRNA data 354 features.

#### 6.4 Integrative clustering using *HC-fused*

The main function for integrative hierarchical clustering is `HC_fused_subtyping()`. The function expects the single-omics data matrices (rows=patients, cols=features) organized as a list. We set the maximum number of possible clusters to `max.k=10`. We use agglomerative clustering based on Ward's method and repeat the calculations 20 times (`HC.iter=20`) to have a good estimate of the contribution of the single-omics to the data fusion process.

```
res <- HC_fused_subtyping(omics = list(mRNA, Methy, miRNA),
                          max.k = 10, this_method = "ward.D",
                          HC.iter = 20)
```

The **res** object is a list containing three elements:

```
names(res)
[1] "cluster" "P"       "S"
```

The vector `cluster` contains the final cluster assignments, `P` is the fused distance matrix and `S` stores information about the contribution of the single data views to the data fusion.

```
cl_fused <- res$cluster
```

```
table(cl_fused)
cl_fused
 1  2  3  4  5  6  7
50 20  2 14 11  4  4
```

*HC-fused* infers seven clusters based on the fused similarity matrix $P$.

#### 6.5 Survival analyses

Finally, the predictive ability of our approach with respect to patient survival is considered.

```
survival <- as.data.frame(clin)
groups <- factor(cl_fused)
names(groups) = rownames(survival)
coxFit <- coxph(Surv(time = Survival, event = Death) ~ groups,
                data = survival, ties="exact")
P_FUSED = round(summary(coxFit)$sctest[3], digits = 40)
P_FUSED
   pvalue
3.232908e-05
```

The resulting $p$-value is highly significant, assuming a significance level of $\alpha = 0.01$.

#### 6.6 Plotting the clustering results

We first assign the patient IDs to the fused distance matrix $P$.

```
colnames(res$P) <- rownames(mRNA)
rownames(res$P) <- rownames(mRNA)
```

A heatmap plot can be generated directly from native R as follows:

```
heatmap(res$P, distfun=as.dist, xlab="patients", ylab="patients",
        cexRow = 0.3 , cexCol = 0.3)
```

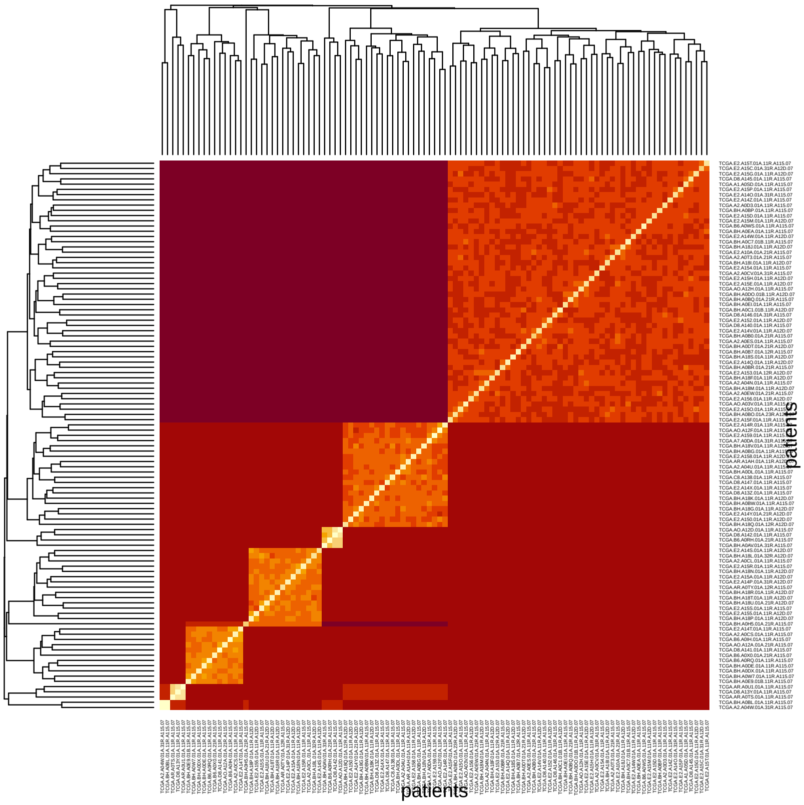


**Supplementary Fig. 8:** Breast cancer data: Clustering results of the fused distance matrix $P$.

The contribution of the single-omics to the clustering can be extracted via the `HC_fused_get_contributions()` function.

```
contrib <- HC_fused_get_contributions(S = res$S ,cluster = cl_fused )
contrib
```

```
          cluster1   cluster2  cluster3  cluster4  cluster5  cluster6  cluster7
omic1   0.10501916 0.11741447 0.3095238 0.1844915 0.1817317 0.1807216 0.3104695
omic2   0.14798154 0.20547533 0.2261905 0.1414435 0.1328039 0.2530103 0.2375532
omic3   0.07160397 0.07631941 0.2142857 0.1106949 0.1118349 0.1626495 0.1839820
omicAND 0.67539532 0.60079079 0.2500000 0.5633702 0.5736295 0.4036186 0.2679953
```

A pie plot may be used to visualize the results.

```
par(mfrow=c(3,3))
pie(contrib[,1], col=c("green","blue","orange","black"),
    xlab="cluster1", labels="")
```

```
pie(contrib[,2], col=c("green","blue","orange","black"),
    xlab="cluster2", labels="")
pie(contrib[,3], col=c("green","blue","orange","black"),
    xlab="cluster3", labels="")
pie(contrib[,4], col=c("green","blue","orange","black"),
    xlab="cluster4", labels="")
pie(contrib[,5], col=c("green","blue","orange","black"),
    xlab="cluster5", labels="")
pie(contrib[,6], col=c("green","blue","orange","black"),
    xlab="cluster6", labels="")
pie(contrib[,7], col=c("green","blue","orange","black"),
    xlab="cluster7", labels="")
plot.new()
legend("center",legend=c("mRNA","Methy","miRNA","ALL"),
        fill=c("green","blue","orange","black"),
        box.lty=0, title="Data␣views", cex=0.75)
```

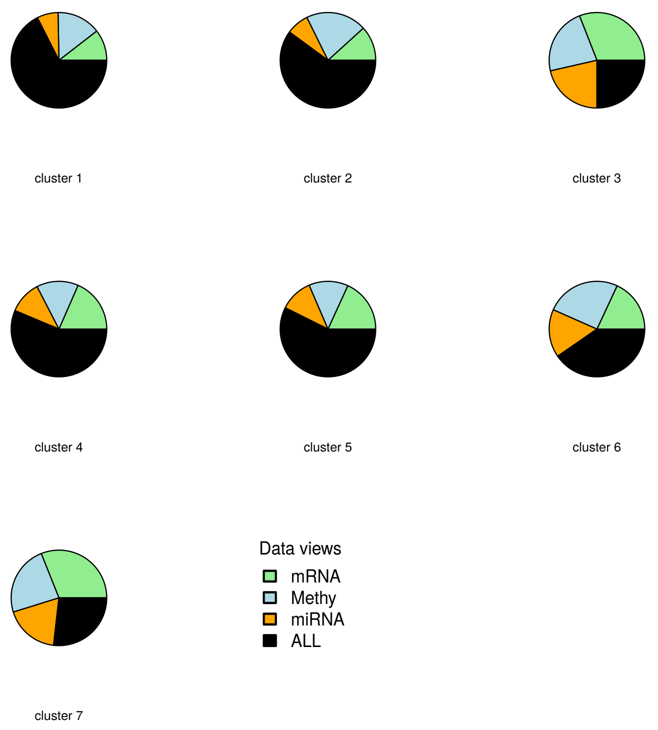


**Supplementary Fig. 9:** Contribution of the views to the hierarchical data fusion. Results are displayed for the TCGA breast cancer data.
